# A General Fluorescence-Based Method for Quantifying and Mapping Biomolecular Polarity In Vitro and In Cells

**DOI:** 10.1101/2023.02.07.526546

**Authors:** Tze Cin Owyong, Riley O’Shea, Mihwa Lee, Jonathan M. White, Paul S. Donnelly, Elizabeth Hinde, Wallace W. H. Wong, Yuning Hong

**Affiliations:** Bio21 Molecular Science and Biotechnology Institute, School of Chemistry, University of Melbourne, Parkville, Victoria 3010, Australia; ARC Centre of Excellence in Exciton Science, University of Melbourne, Parkville, Victoria 3010, Australia; Department of Biochemistry and Chemistry, La Trobe Institute for Molecular Science, La Trobe University, Melbourne, VIC 3086 Australia; School of Physics, University of Melbourne, Parkville, Victoria 3010, Australia

## Abstract

Spatial discretization of biomolecules in the complex cellular environment is crucial for biomolecular form and function. The ability to better understand the driving force of spatial discretization of biomolecules in the complex cellular matrix remains a challenging task. We report on the robust polarity sensitive solvatochromic probe, **FLAM**, in conjunction with spectral phasor analysis as a general method for studying environmental polarity in biological systems. We find that phase separated proteins of SFPQ have distinct polarity depending on the type of phase separation occurring, suggesting that polarity plays a role in the formation of phase separated condensates. When using **FLAM** in cells, distinct subcellular environmental polarity distribution but similar trend of changes is observed for cells under similar type of stressors. Taken together, our method puts forth an exciting development in the tool set for the study of phase separation.

## Introduction

The intracellular environment is complex and heterogenous, with a multitude of biological processes concurrently occurring in distinct chemical environments. Organisation and control over these biological processes can be achieved by spatial discretisation, with diffusion of molecules in these compartments playing a further role in these processes. Many compartmentalised organelles rely on a membrane for separation, with some examples including the nucleus, mitochondria and endoplasmic reticulum (ER), among others. On the other hand, there are many membraneless organelles in various parts of the cell, including Cajal bodies, paraspeckles in the nucleus as well as P-bodies and stress granules in the cytoplasm. Each of these membraneless organelles contribute towards or perform specific pathways or functions.(*1*) An example is the formation or presence of ribonucleoprotein granules (e.g. stress granules or P-bodies), that are crucial in the regulation of mRNA translation and contribute to the proteostasis machinery.(*2*)

Current strategies to study the properties of these transient biomolecular condensates include a range of experimental techniques such as fluorescence recovery after photobleaching (FRAP), fluorescence correlation spectroscopy (FCS), Förster resonance energy transfer (FRET), single particle tracking (SPT) and fluorescence lifetime imaging microscopy (FLIM). Specific information such as transport and rearrangement, concentration and distances of molecules can be obtained by using these techniques, contributing to the understanding of the biophysical properties and functions of these membraneless organelles. Extensive mutagenesis identifies the molecular grammar of amino acid sequence governing the formation of membraneless organelles.(*3*) These observations imply the importance of the physiochemical properties of the biomolecules and their surrounding environment, such as viscosity, polarity, macromolecular crowding, and ionic strength, as the driving forces of phase separation.(*4–7*) The ability to estimate environmental polarity and other physical factors within a heterogeneous condensate would thus provide useful information in elucidating the formation, interaction, and nature of various condensates. A viable strategy to address these questions is the application of specialised imaging techniques and analysis in combination with environmental sensitive fluorescent probes.

Spectral phasor analysis allows us to deconvolute and identify the emission spectra of individual pixels in a spectral image on a global scale. In previous work, we demonstrated that acquisition of spectral images followed by spectral phasor analysis enabled the estimation of the intracellular local environment of a bound thiol reactive dye, **NTPAN-MI**. Upon proteostasis stress, the estimated dielectric constant in cells became more heterogeneous.(*8*) To better understand this observed phenomenon as well as demonstrate the broad applicability of spectral phasor analysis, herein, we adopt a non-covalent based approach with a solvatochromic fluorophore, **FLAM**, which bears good solvatochromic properties, cell permeability and compatibility, to quantify polarity and its shifts *in vitro* and in cell (Fig. 1). Such a non-covalent dye could allow us to apply the method universally in purified biomolecules *in vitro* and in various cellular compartments, membraneless organelles and condensates in cells undergoing proteostasis stress.

**Fig. 1.**
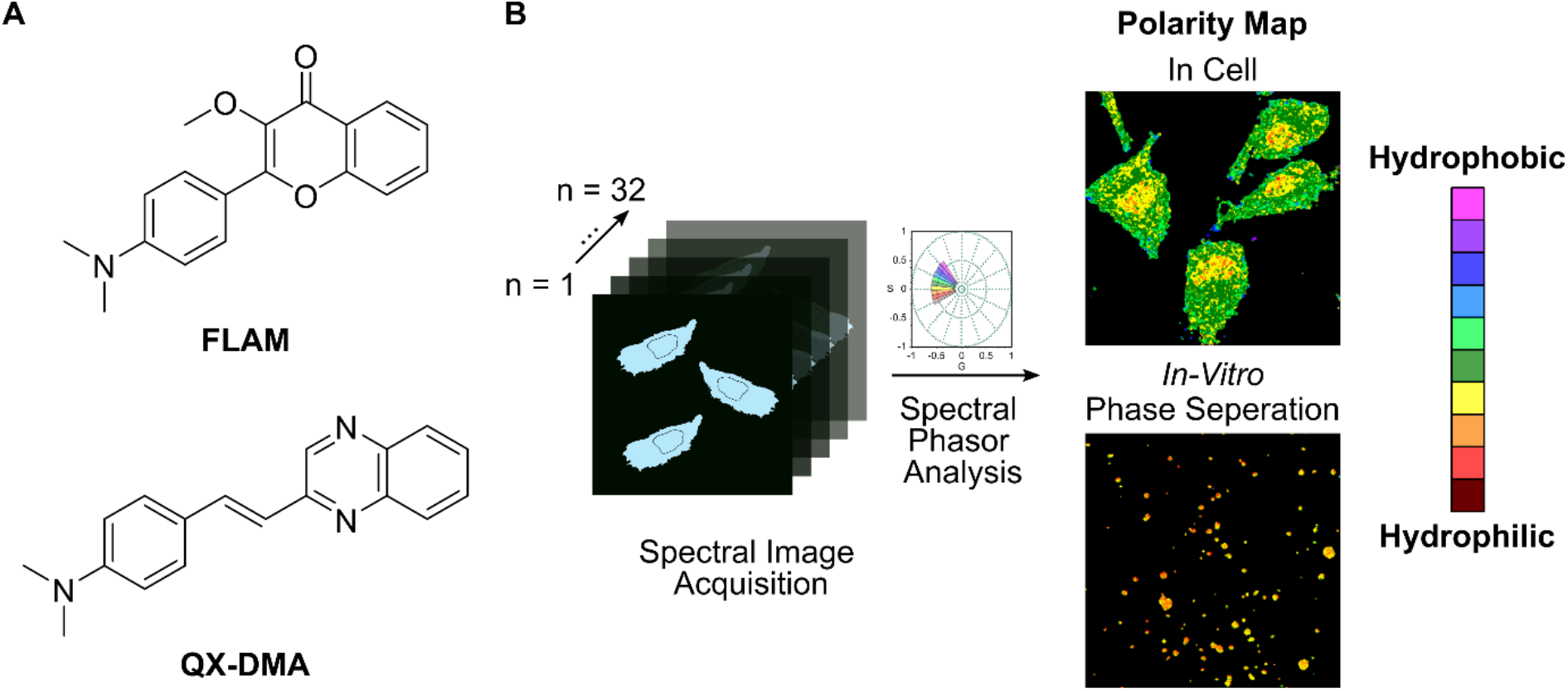
Solvatochromic fluorophores coupled with image analysis enables estimation of polarity. (**A**) Solvatochromic dyes synthesized for this study. (**B**) Schematic overview of experiments for estimation of environmental polarity in biological system.

## Results

### Synthesis & photophysical characterisation of polarity sensitive probes

Various strategies can be used to design a polarity sensitive probes, such as using charge transfer, intramolecular proton transfer, conformational changes, isomerisation, and aggregation.(*9*) In this work, we designed two solvatochromic fluorophores based on their electron push-pull structure facilitating intramolecular charge transfer. We first conducted in silico verification of our hypothesis. Calculated HOMO and LUMO orbitals for **QX-DMA** and **FLAM** showed that the HOMO was more localised on the electron rich dimethylamine ring while the LUMO was centred on electron withdrawing groups of each respective molecule (Fig. 2A). The relative discrete HOMO and LUMO orbitals for both molecules should give rise to an intramolecular charge transfer state and subsequent solvatochromic photoluminescent behaviour upon photoexcitation. **QX-DMA** was also seen to have a smaller HOMO-LUMO energy gap of 2.92 eV as compared to **FLAM** of 3.37 eV.

**Fig. 2.**
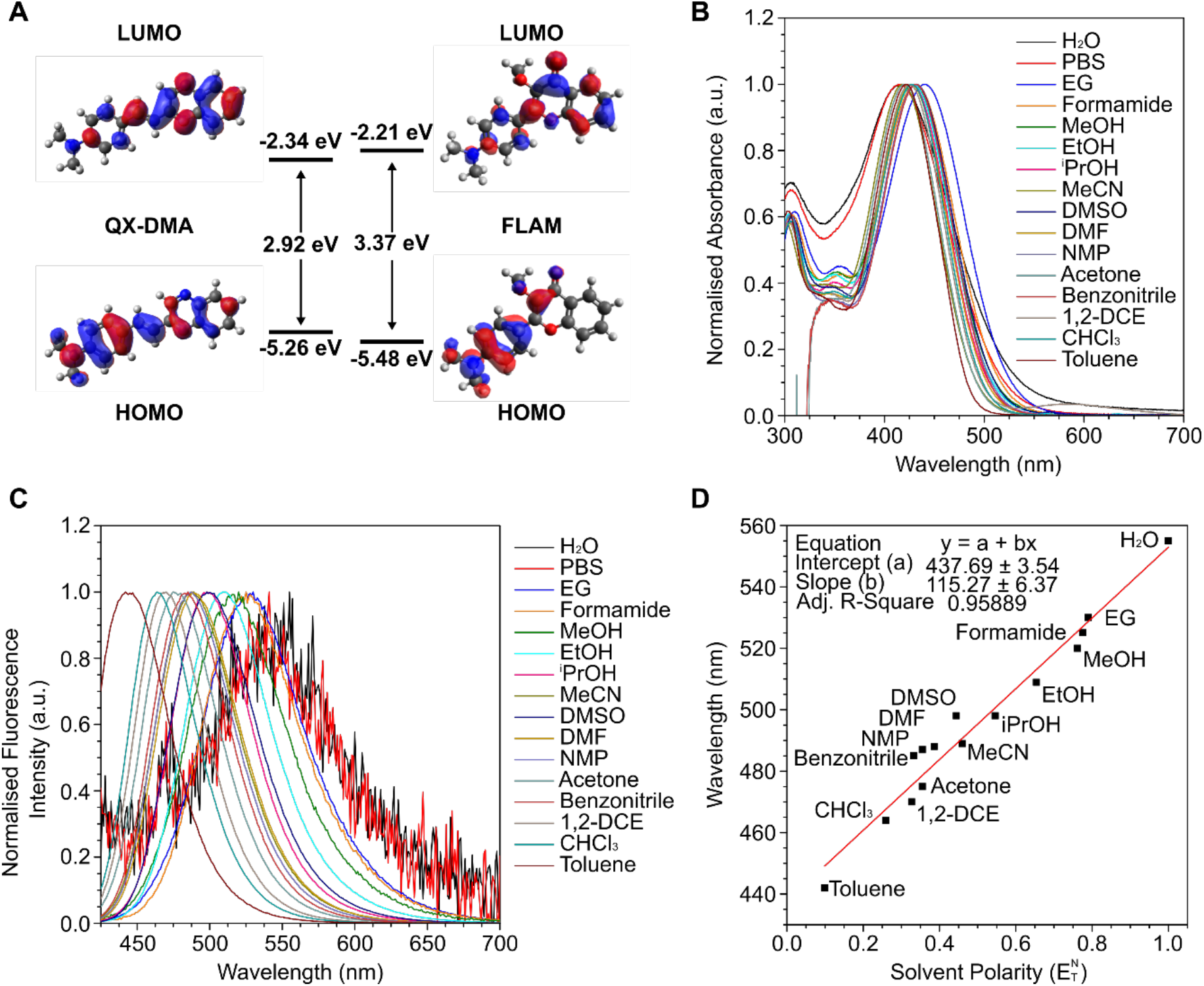
Photophysical characterisation of solvatochromic behaviour of dyes. (**A**) DFT calculations for environment sensitive dyes, **QX-DMA** and **FLAM**. DFT calculations were carried out at the B3LYP/6-31+(d,p) level with IEFPCM model in water. No negative frequencies for optimised structures were observed. (**B**) Normalised absorbance spectra of **FLAM** in different solvents. (**C)** Normalised fluorescence emission spectra of **FLAM** in different solvents. (**D**) Relationship of measured fluorescence emission maxima wavelength and solvent polarity for **FLAM**. Solvent polarity was based on reported values, with PBS excluded from linear plot (see **D**).(*18*) 405 nm excitation wavelength was used for fluorescence emission spectra measurements. PBS was Na_2_HPO_4_ at 20 mM concentration. 10 μM dye concentration was used for all measurements. 1,2-DCE: 1,2-dichloroethane; iPrOH: isopropanol; EG: ethylene glycol.

We next performed multi-step synthesis and purification of the two fluorophores. For **QX-DMA**, the strong electron acceptor, 2-methylquinoxaline, was synthesised by condensation of pyruvaldehyde and 1,2-phenylenediamine in good yields. Subsequently, chlorination with triisocyanuric acid (TCCA), Arbuzov reaction with triethylphosphite and Horner-Wadsworth-Emmons (HWE) reaction with *p*-dimethylaminobenzaldehyde gave the final product in fair yield over 3 steps. For **FLAM**, an aldol condensation of 4-dimethylaminobenzaldehyde and 2-hydroxyacetophenone followed by cyclisation with hydrogen peroxide gave the flavone core in moderate yield over 2 steps. Subsequent alkylation of free hydroxy group gave solvatochromic probe **FLAM** in good yield (Scheme S1 in the Supporting Information). The intermediates and final products were characterised using ^1^H, ^13^C NMR spectroscopy and HRMS. X-ray diffraction crystallography further confirmed the structures of the final products **QX-DMA** and **FLAM** (Figure S1 & Table S1, 2).

To characterize their solvatochromism, we measured their absorption and emission spectra in different solvents. The selection of a wide range of solvents with varying polarity 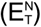, including more physiologically relevant polarities enabled the relationship with the fluorescence emission maxima to be established. First, both **QX-DMA** and **FLAM** showed minimal changes in absorbance, with an absorbance maximum centred around 430 and 390 nm respectively (Fig. 2B, S2A, and S2C). Fluorescence emission spectra, on the other hand, showed that both **QX-DMA** and **FLAM** exhibited a strong solvatochromic behaviour, showing positive solvatochromism with emission maximum shifting over 100 nm from non-polar to polar solvents (Fig. 2C, S2B and S2D). This behaviour is characteristic of a photoinduced charge transfer mechanism. Upon excitation by light, a polarity sensitive fluorophore undergoes an intramolecular charge transfer that involves redistribution of electrons within the molecule, resulting in a fast change in the dipole moment of the molecule. The surrounding solvent molecules can reorganise to create a relaxed state of minimum energy. As such, solvents with higher polarity, such as DMSO, will cause a larger red shift in the emission spectra of a polarity sensitive dye due to the decrease in energy of the relaxed state. Comparison between **QX-DMA** and **FLAM** showed that only **FLAM** had a strong linear relationship for fluorescence emission maxima and solvent polarity (Fig. 2D and S3). Subsequent experiments and applications were then focused on **FLAM**.

To further validate the dyes for use in cells, we examined the effect of pH and viscosity of the two dyes at physiologically relevant conditions. Minimal changes were observed within physiological pH range (Fig. S4). The emergence of a new peak at shorter wavelength was observed only at very low pH when the dimethylamino unit was protonated. Despite the solvatochromic nature of **FLAM**, we observed a less profound change of fluorescence intensity in response to increased viscosity, suggesting minimal effects of restriction of intramolecular rotation on **FLAM** (Fig. S5). Altogether, these results suggest that **FLAM** could be a good candidate fluorophore for reporting on polarity for our subsequent application in vitro and in cells.

### FLAM measures polarity of biomolecules and in phase separated protein droplets

Since the emission wavelength of **FLAM** is dependent on the solvent polarity, we asked whether we can make use of this property to measure the polarity of the biomolecules surrounding **FLAM**. We chose two model proteins, bovine serum albumin (BSA) and *β*-Lactoglobulin (*β*-Lac), to compare with highly charged biomolecules such as DNA and RNA. Consistent with our hypothesis, incubation of **FLAM** with both BSA and *β*-Lac in HEPES buffer gave a fluorescence spectrum in the blue region, while the addition of DNA and RNA led to a significant redshift of the spectrum (Fig. 3A). Interestingly, the estimated polarity of these model proteins from the spectra was similar to acetonitrile while those of the DNA and RNA were close to ethanol and methanol respectively (Fig. 3A).

**Fig. 3.**
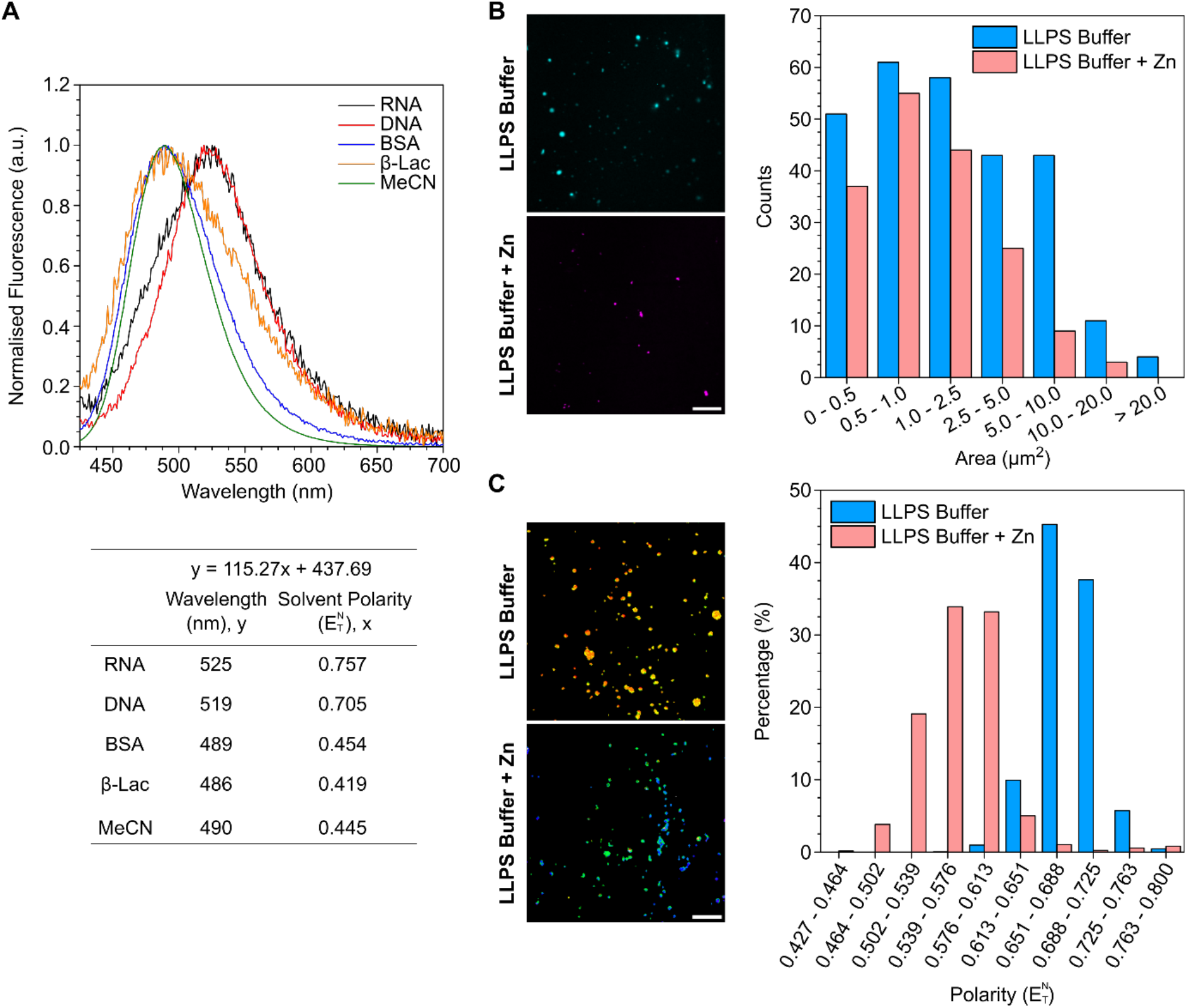
*In-Vitro* characterisation of FLAM with biomolecules and protein phase separation studies. (**A**) Normalised fluorescence emission spectra for **FLAM** in the presence of biomolecules. 100 μg/mL of the respective biomolecules was incubated with 10 μM **FLAM** and 20 mM HEPES were used for all measurements. (**B**) Representative confocal microscopy images of phase separated protein droplets, Ex/Em 405/490 – 560 nm channel of **FLAM** shown, and quantification of phase separated droplet size and count. (**C**) Representative confocal microscopy images of protein droplet polarity resolved with spectral phasor analysis and quantification of environmental polarity distribution in phase separated droplets. 32 channel spectral acquisition was obtained with 405 nm excitation; 420 – 580 nm emission with 5 nm bin widths. For protein phase separation experiments, LLPS buffer-treated SFPQ protein gave liquid-like phase separated droplets. In the presence of zinc (Zn), SFPQ protein gave solid-like condendates. The final composition of the LLPS buffer inducing phase separation was 20 mM HEPES (pH 7.4), 150 mM KCl, 60 mM L-Arg, 1.5% glycerol, while that of the zinc buffer was 20 mM HEPES (pH 7.4), 150 mM KCl, 60 mM L-Arg, 1.5% glycerol, 3 μM ZnCl_2_. 0.8 mg/mL SFPQ ΔN protein (residues 276 – 706) was used with 5 μM **FLAM** for all conditions. n = 3 images, Scale bar: 20 μM.

These results demonstrated the capability of using **FLAM** to quantify biomolecular polarity in solution. We next asked whether we can use **FLAM** to measure the polarity within phase separated droplets. Splicing factor proline- and glutamine-rich (SFPQ) protein, an RNA- and DNA-binding protein involved in DNA repair and paraspeckle formation, was chosen for this study.(*10*) SFPQ has been shown to form liquid-like droplets with buffer containing physiological salt concentration, or aggregate in the presence of zinc ions.(*11, 12*) We carried out confocal microscopy experiments incubating **FLAM** and SFPQ in the two conditions to image the respective liquid-droplets and solid-like condensates. First, we observed that **FLAM** was localised with both types of SFPQ condensates, with negligible background fluorescence intensity. Due to the low background, we could then use an intensity based strategy to segment and quantify the number and size of phase separated condensates (Fig. 3B). Upon confirming the feasibility of using **FLAM** to stain for and quantify SFPQ condensates, we moved on to characterise the environment polarity of SFPQ condensates using spectral phasor analysis. We observed that the liquid-like condensates from the buffer containing physiological salt concentration gave a drastically more hydrophilic environment as compared to the solid-like condensates with zinc treatment (Fig. 3C). This is in line with previous reports that hydrophobicity and multivalent interactions contribute to solid-like protein aggregation and liquid-liquid phase separation (LLPS) respectively.(*13–16*)

### Evaluation of FLAM for cell imaging experiments

After validating the feasibility of using **FLAM** to estimate biomolecular polarity in vitro, we next moved on to *in cellulo* experiments. Cytotoxicity of **FLAM** was evaluated in both HeLa and A549 cell lines, showing excellent cell viability for 30 min staining time across a wide range of dye concentration. Cell viability remained above 80% over 24 and 48 h incubation with dye concentration up to 50 and 10 μM respectively (Fig. S6). For our staining experiment, we chose 5 μM for 0.5 h, which is sufficient for FLAM to penetrate through the cell membrane and enable fluorescence based experiments. The cell cytotoxicity experiments demonstrated that **FLAM** had low cytotoxic effects on cells and could be used for longer term monitoring of cells.

Following confirmation of low cytotoxic effects for **FLAM**, we then examined its cell uptake and intracellular localisation. To be able to use counterstains, dye crosstalk experiments showed minimal signal bleedthrough for each dye and demonstrated good spectral compatibility of **FLAM** with a range of commercially available fluorescent dyes with varying spectral characteristics (Fig. S7). Dye localisation studies were then carried out with HeLa and A549 cell lines. Good uptake of **FLAM** was observed for both cell lines with 0.5 h dye staining at 37 ^°^C. Inside the cells, **FLAM** distributed across both cytoplasm and nucleus with enhanced fluorescence colocalised with ER (Fig. 4A). When we collected fluorescence signals at a shorter (410-480 nm) and a longer (490-560 nm) wavelength region, corresponding to the low and high polarity environment, we observed a different staining pattern in particular for the cytoplasmic region. This staining pattern remained the same in both cell lines, suggesting that **FLAM** could be applied as a universal non-covalent environment polarity probe, across cell lines. A z-stack microscopy experiment was then carried out on HeLa cells. Inspection of the 3-dimensional staining pattern of **FLAM**, with co-stains, further confirmed cellular localisation and uptake in both cytoplasmic and nuclear compartments (Fig. S8).

**Fig. 4.**
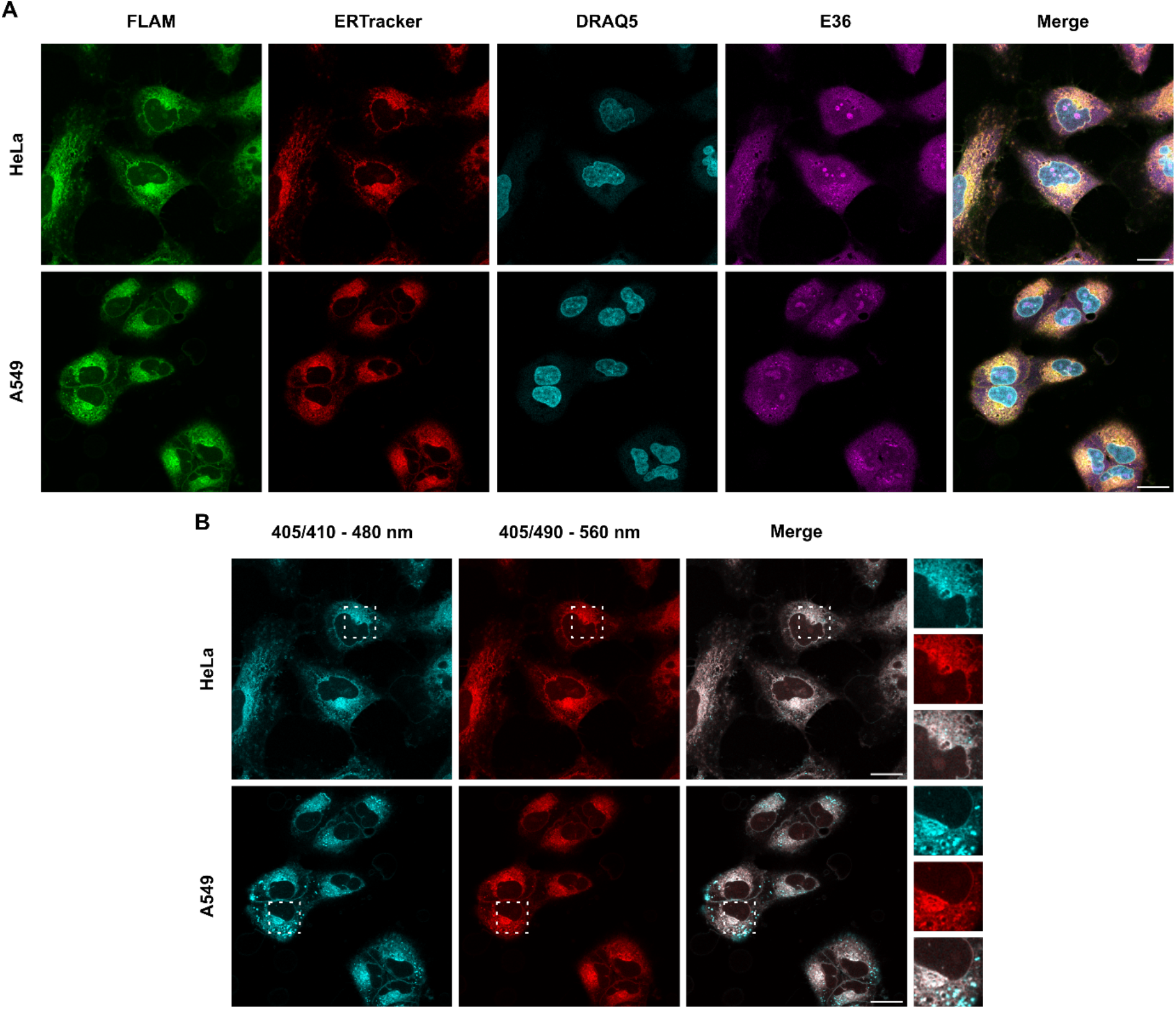
Cellular localisation analysis for FLAM. (**A**) Colocalisation imaging experiments with ERTracker™ Red, DRAQ5™ and E36 stains were used to visualise endoplasmic reticulum, nucleus (DNA) and RNA respectively. Cells were stained with 5 μM **FLAM**, 1 μM DRAQ™ and E36, and 500 nM ERTracker™ Red. 405/490 – 560 nm channel of **FLAM** shown. (**B**) Differences in cell staining localisation at different emission channels. Scale bar, 20 μm.

We next evaluated the photostability of **FLAM**. Photostability of fluorophores is particularly important for their sensitivity and reliability for quantitation in biological systems. Continuous acquisition for a set of 50 images of **FLAM** stained HeLa cells was conducted and the changes of fluorescence intensity were subsequently analysed (Fig. S9). Excellent dye resistance to photobleaching was observed, with 88% fluorescence intensity retained after the long acquisition, demonstrating good photostability of **FLAM** and providing us confidence of using this dye for further imaging experiments requiring multiple acquisitions over time.

### FLAM reveals subcellular polarity change in cells under different stresses

Following the characterisation of dye behaviour, we confirmed that **FLAM** has excellent biocompatibility, cell uptake, and resistance against photobleaching. Such traits, in combination with its positive solvatochromism enabled acquisition of spectral images for cells and to apply spectral phasor analysis to estimate the general intracellular polarity environment based on emission of **FLAM**. We first selected a series of treatment conditions, which we generally summarized into oxidative, osmotic, transport pathway and protein unfolding stress to study the polarity changes in response to such conditions. Following acquisition and spectral phasor analysis, we observed that cells had different polarity environments, dependent on the treatment conditions (Fig. 5A). An example can be found when comparing control, arsenite and H_2_O_2_treatments, between those two oxidative stressors, the H_2_O_2_treatment resulted in a more hydrophobic environment compared to arsenite treatment. We then quantified our observations by analysing the pixel population of each polarity range for individual cells in their respective treatment conditions.

**Fig. 5.**
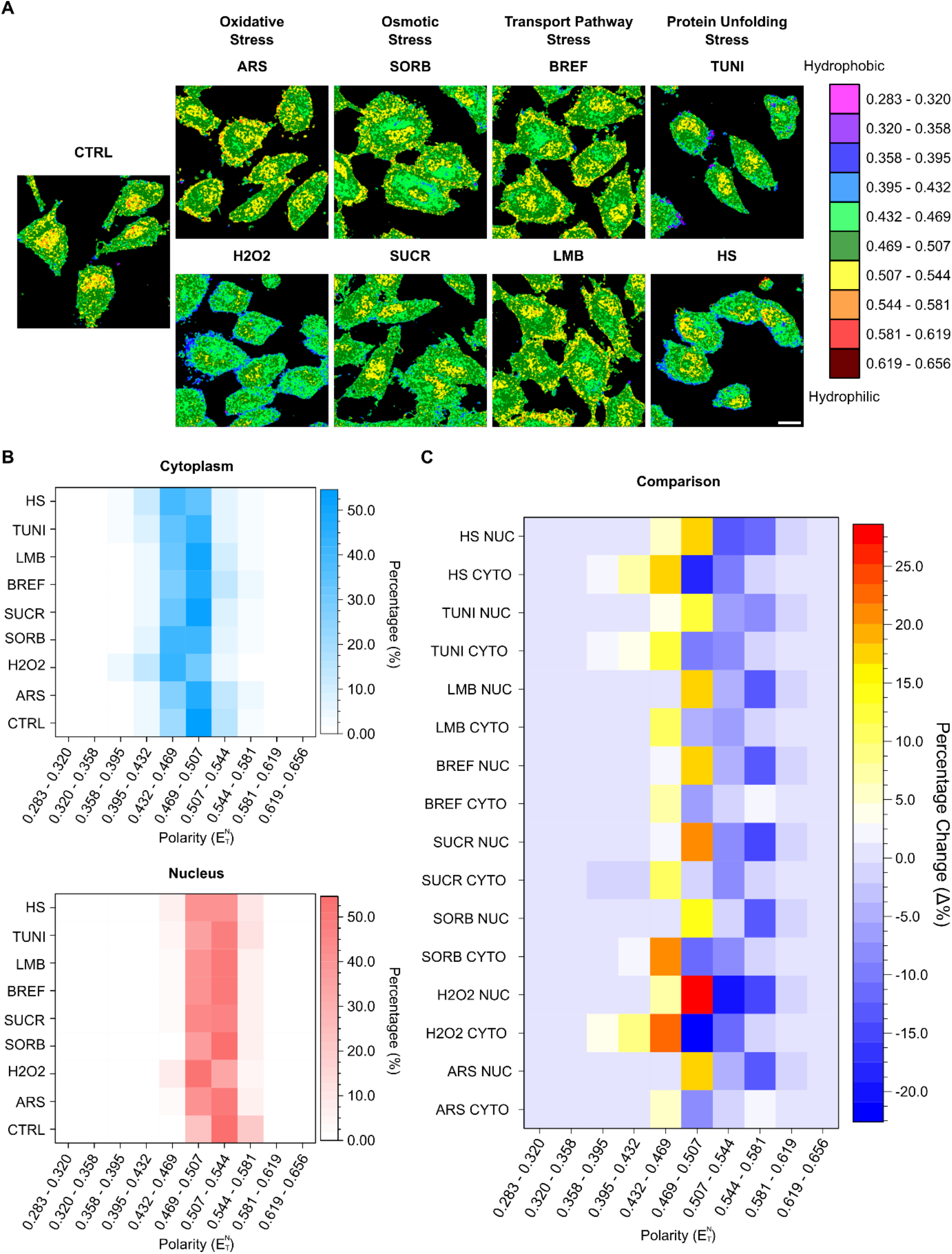
Cellular environmental polarity estimation using spectral phasor analysis of FLAM fluorescence. (**A**) Representative confocal microscopy mapping of intracellular polarity for untreated and treated cells stained by **FLAM** resolved with spectral phasor analysis. 32 channel spectral acquisition was obtained with 405 nm excitation; 420 – 580 nm emission with 5 nm bin widths. Scale bar: 20 μM. (**B**) Distribution of environmental polarity in cytoplasm and nucleus for untreated and treated HeLa cells. 32 channel spectral acquisition was obtained with 405 nm excitation; 420 – 580 nm emission with 5 nm bin widths. Spectral phasor analysis was based on 10 images, n = 2 biological replicates of 5 images, with a total of 37 – 58 cells analysed for each treatment condition. (**C**) Change of cellular environmental polarity, obtained by spectral phasor analysis, for respective cellular compartments, of cytoplasm and nucleus, and comparison between treated against untreated cells. CTRL: control; H2O2: hydrogen peroxide (H_2_O_2_); SORB: sorbitol; SUCR: sucrose; BREF: brefeldin A; LMB: leptomycin B; TUNI: tunicamycin; HS: heat shock.

To better understand the heterogeneity of the intracellular polarity, we segmented the cells generally into cytoplasm and nucleus based on the fluorescence intensity (Fig. S10). Upon creating cytoplasm and nucleus masks, single cell masks of respective cellular compartments were acquired, to then allow for selection of our cellular region of interest in spectral images and subsequent spectral phasor analysis (Fig. S10). Upon analysis of 37-58 cells, we found that the cytoplasm consistently had a more hydrophobic environment than nucleus in general, with the polarity centred around 0.432 – 0.507 and 0.469 – 0.544 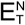 for cytoplasm and nucleus respectively (Fig. 5B). By comparing the polarity change for treated against untreated cells and their respective cellular compartments (e.g. arsenite cytoplasm against control cytoplasm), we can see that cells under stress had an overall increase in hydrophobic environments for both cytoplasm and nucleus, with the degree of polarity shifts dependent on the treatment condition (Fig. 5B and 5C). Furthermore, we observed different degrees of changes in localisation and distribution of environmental polarity, depending on treatment conditions (Fig. 5C). In the case of heat shock, tunicamycin and H_2_O_2_, a significant shift to a much lower polarity, hydrophobic environment was observed in cytoplasm compared to that of control. Fig. 5C also revealed that similar type of stressors, such as LMB and Brefeldin A treatment, both interfering with the transport pathways, led to very similar shift in the intracellular polarity, highlighting the robustness of this method.

## Discussion

In this work, we have designed and synthesized two polarity sensitive fluorophore, **QX-DMA** and **FLAM**, based on the electron push-pull structural configuration. **FLAM** displayed an excellent linear relationship of emission wavelength and solvent polarity index 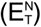 enabling estimation of polarity in biological systems and thus was used for subsequent experiments. We first demonstrated that the emission spectral profile of **FLAM** is dependent on the type of biomolecules in solution, showing proteins are generally more hydrophobic than the highly charged nucleic acids with RNA being even more hydrophilic than DNA. The polarity of these biomolecules was found within the polarity range of DMSO/acetonitrile and methanol, which is consistent with previously reported values.(*7, 17*)

Spectral phasor analysis allows us to deconvolute and identify the emission spectra of individual pixels on a global scale regardless of the intensity. This eliminates the variation of local dye concentrations. By harnessing spectral phasor analysis, we successfully applied **FLAM** to visualize and quantify two different types of condensates formed by the RNA binding protein, SFPQ. The very low background fluorescence enables intensity-based quantification of size and count of the phase separating droplets with confocal microscopy. Further applying spectral phasor analysis enabled the characterization of environmental polarity within both liquid- and solid-like condensates. A distinct polarity in those two types of SFPQ condensates demonstrates that hydrophobic and multivalent interactions play a critical role in driving the formation of liquid- and solid-like condensates respectively, supporting the molecular grammar controlling the formation of those two types of biocondensates once concentration threshold is reached.

With excellent cell compatibility, uptake, retention and photostability, **FLAM** was applied to quantify cellular environmental polarity in cells under different stress treatments. Due to the complexity of the cellular environment, such as the coexistence of proteins, nucleic acids and other molecules, a smaller range of polarity distribution was observed in cells compared to in vitro with a single type of biomolecule only. Within the small polarity range, we can still observe different degrees of polarity shift in segmented nucleus and cytoplasm of cells under different treatments, providing information on the role of polarity in cellular proteostasis regulation and stress response. Overall, we show the importance of environmental polarity in biological systems and put forward a robust fluorophore coupled with analysis technique to reliably probe polarity in situ.

## Acknowledgements

We thank BOMP, University of Melbourne for access to the confocal microscope and Bio21 Mass Spectrometry and Proteomics Facility for technical support and access to mass spectrometers. This work was supported by grants to Y.H. (Australian Research Council FT210100271 and National Health and Medical Research Council APP1101803) and WWHW (Australian Research Council CE170100026).

