## Supporting information for "A General Fluorescence-Based Method for Quantifying and Mapping Biomolecular Polarity In Vitro and In Cells"

### Supplementary Information

#### 1 Materials and Methods

##### Materials

##### Synthesis Characterisation

$^{1}H$ (400MHz or 500MHz) & $^{13}C$ NMR (100MHz, 125MHz) spectra were acquired on Agilent MR400 or Agilent DD2 instrument. The chemical shift data for each signal are given as δ. High-resolution mass spectra were acquired using a Thermo Scientific Q Exactive Plus Orbitrap LC-MS/MS instrument.

##### X-ray Diffraction Experiments

X-ray diffraction intensity data for **QX-DMA** and **FLAM** were collected with a Rigaku Synergy Diffractometer using either Cu – Kα radiation with the temperature during data collection maintained at 100.0(1) K using an Oxford Cryosystems cooling device. The structures were solved by direct methods and difference Fourier synthesis. Thermal ellipsoid plots were generated using the program Mercury integrated within the WINGX suite of programs.

##### Photophysical Characterisation

Absorbance and fluorescence spectra were obtained on a Cary 300 UV-Vis spectrometer and Cary Eclipse fluorimeter (Agilent Technologies Inc., Santa Clara, CA, USA), respectively. Data were plotted using Origin 2018 (OriginLab Corp., Northampton, MA, USA).

##### DFT Calculations

DFT calculations were carried out using B3LYP/6-31+g(d,p) level with IEFPCM model in water with Gaussian16, using the University of Melbourne HPC system, Spartan, for all calculations. Visualisation of structures were done using Avogadro 1.20 and GaussView 5.0. No negative frequencies for optimised structures were observed.

**Cell Culture**

HeLa and A549 cells were cultured in DMEM (Life Technologies, Catalog Number: 11965118) supplemented with 10% fetal bovine serum (Corning, Australia origin, Catolog Number: 35-076-CV) at 37 °C in 5% $CO_2$ air with humidification.

**Cell Viability**

2 x $10^3$ cells were plated onto 96-well plates 24 h prior to dye application. FLUOstar Omega plate reader (BMG Labtech) in fluorescence intensity mode with excitation at 544 nm and emission at 590/10 nm was used to assess cell viability using Alamar Blue assay (Thermo Fisher Scientific, Catalog number: DAL1025).

**Cell Stress and Staining**

**FLAM** was dissolved in DMSO as a 5 mM stock solution and were kept at -20 ºC in the dark. 1.5 x $10^4$ HeLa or A549 cells were plated on an ibidi µSlide 8 Well, ibiTreat (ibidi, Catalog number: 80826-90) or 7.0 x $10^4$ on an ibidi Glass Bottom Dish 35mm (ibidi, Catalog number 81158) for fixed cells. Proteostasis stress inducing drugs were added to plated HeLa cells before treatment with dye. Plated cells were treated with freshly diluted dye (5 µM) for 30 min at 37ºC and subsequently washed. Cells were fixed on plate with 4% (w/v) paraformaldehyde (PFA) in PBS for 15 min at room temperature.

**Cell Stress Conditions**

Sodium Arsenite: 1 h treatment with 500 µM; $H_2O_2$: 1 h treatment with 200 µM; Sorbitol: 1 h treatment with 400 mM; Sucrose: 1 h treatment with 200 mM; Brefeldin A: 1 h treatment with 10 µg/mL; Leptomycin B: 30 min treatment with 5 nM; Tunicamycin: 8 h treatment with 5 µg/mL; Heat Shock: incubated at 42 °C for 30 min.

**Confocal Laser Scanning Microscopy**

After staining, cells were fixed with 4% (w/v) PFA in PBS for 15 min at room temperature. Images were acquired on a Zeiss LSM 880 microscope using a 63x objective lens. For image acquisition, the pixel frame size was set at 512x512 and the pixel dwell time was 32.7 µs. ERTracker™ Red, DRAQ5™ and

E36 stains were used to visualise endoplasmic reticulum, nucleus (DNA) and RNA respectively. Cells were stained with 5 μM **FLAM** (excitation: 405 nm; emission 410 – 470, 490 – 560 nm), 1 μM DRAQ$^{TM}$ (excitation: 633 nm; emission: 680 – 740 nm) and E36 (excitation: 488; emission 490 – 560 nm), and 500 nM ERTracker$^{TM}$ Red (excitation: 561; emission (565 – 625 nm).

For spectral image acquisition, cells were stained with 5 μM **FLAM** the pixel frame size was set at 256x256 and the pixel dwell time was 16.4 μs. 405 nm excitation laser was used, with emission taken from 420 – 580 nm gated into 32 channels each with 5.0 nm bin width. Image quantification was carried out using Fiji (ImageJ) and SimFCS software.

***In-Vitro* Phase Separation Assays**

SFPQ ∆N protein (residues 276 – 707) was expressed and purified for *in-vitro* experiments according to previously reported procedure with some modifications. To keep the protein from phase separation, 0.2 M L-Arg was included in the purification buffer. The final protein was in 20 mM HEPS (pH 7.4), 0.5 M KCl, 0.2 M L-Arg, 5% glycerol. To induce phase separation, the protein sample was diluted 3/10 by adding no salt buffer (20 mM HEPES, 1 mM DTT, pH 7.4) with or without 3 μM zinc chloride, to give a final solution volume of 10 μL containing 20 mM HEPES (pH 7.4), 150 mM KCl, 60 mM L-Arg, 1.5% glycerol or final zinc buffer of 20 mM HEPES (pH 7.4), 150 mM KCl, 60 mM L-Arg, 1.5% glycerol, 3 μM $ZnCl_2$. and a protein concentration of 0.80 mg/mL. No salt buffer was added rapidly but carefully to avoid bubbles and samples were flick mixed immediately after dilution. For negative controls, storage buffer (20 mM HEPS (pH 7.4), 0.5 M KCl, 0.2 M L-Arg, 5% glycerol) was added to samples instead of no salt buffer.

#### 2 Synthesis Procedures and Characterisation

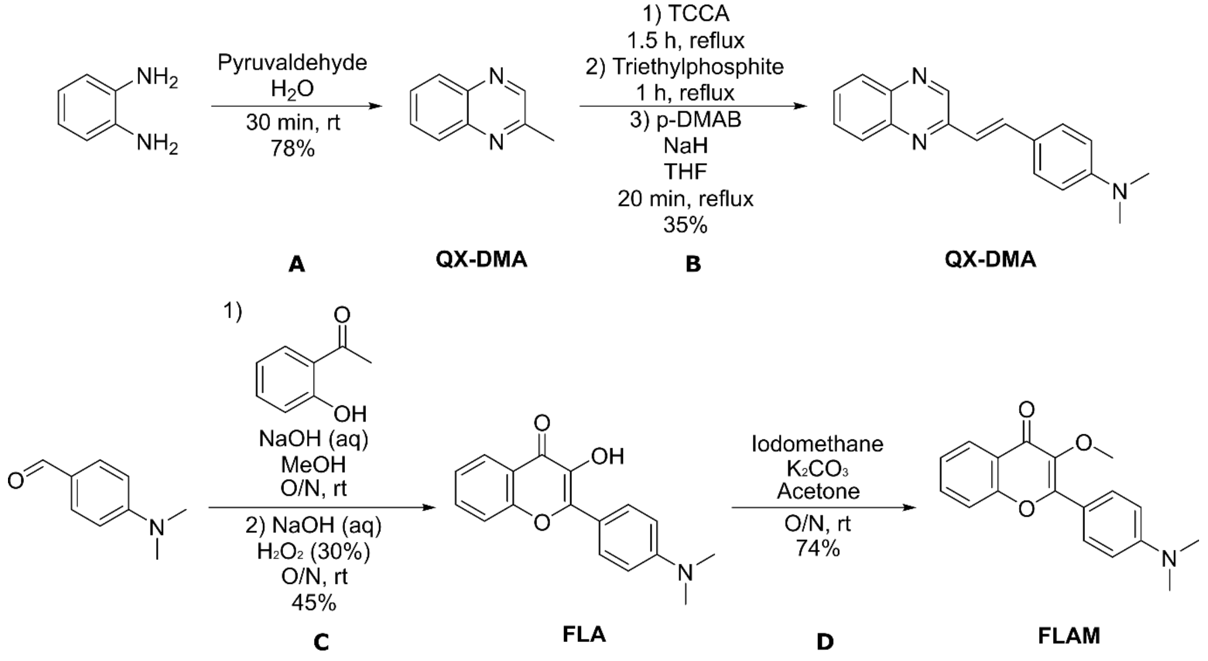


**Scheme S1**: Synthesis scheme for the formation of solvatochromic probe **QX-DMA** and **FLAM**. (A) Formation of electron acceptor core. (B) Chlorination with TCCA, followed by Arbuzov reaction with triethylphosphite and HWE reaction. (C) Aldol condensation followed by cyclisation. (D) Alkylation reaction.

**QX**

1,2-phenylenediamine (10.0 g, 92.5 mmol) was added portionwise into a stirring mixture of pyruvaldehyde (6.65 g, 92.5 mmol) and water (150 mL). The resulting reaction mixture was then stirred at room temperature for 30 min. Upon completion, the reaction mixture was extracted with DCM (x3) and washed water (x3) before being dried over magnesium sulfate, filtered through a short plug of silica and concentrated to give the pure product as a dark yellow oil in 78% yield (10.4 g). $^1$H NMR (500 MHz, $CDCl_3$) δ 8.75 (s, 1H), 8.07 (d, *J* = 8.2 Hz, 1H), 8.01 (d, *J* = 8.1 Hz, 1H), 7.76-7.69 (m, 2H), 2.78 (s, 3H). $^{13}$C NMR (125 MHz, $CDCl_3$) δ 153.78, 146.01, 142.06, 140.95, 130.01, 129.16, 128.92, 128.64, 22.60. HRMS (ESI+): *m*/*z*. 145.07603 [$C_9H_9N_2$ $(M+H)^+$, calcd 145.07602].

**QX-DMA**

2-Methylquinoxiline (500 mg, 3.47 mmol) was added in 2 mL of chloroform, stirred and heated at reflux under $N_2$. To this mixture, trichloroisocyanuric acid (350 mg, 1.53 mmol) was added in portions over 90 min. Upon completion, the reaction mixture was then cooled to room temperature and poured into 5 mL of water, extracted with chloroform (5 mL), washed with saturated aqueous sodium bicarbonate before being dried over magnesium sulfate, filtered and concentrated to give a pale yellow solid.

Without further purification, 2-(chloromethyl)quinoxaline was added to triethylphosphite (3.62 mL) and stirred at reflux for 1 hour. Upon completion, the reaction mixture was then cooled to room temperature before the excess triethylphosphite was removed via distillation under vacuum.

Without further purification, diethyl (quinoxalin-2-ylmethyl)phosphonate (3.47 mmol) was added in THF (3 mL) and added dropwise to a solution of sodium hydride (87.9 mg, 3.82 mmol) in THF (3 mL) at room temperature under $N_2$. A solution 4-(dimethylamino)benzaldehyde (517 mg, 3.57 mmol) in THF (4 mL) was added to the reaction mixture and was stirred and heated at reflux for 20 minutes. Upon completion, the reaction mixture was then cooled to room temperature and quenched with water. The organic layer was extracted with $CHCl_3$ (x3), washed with water (x3), dried over magnesium sulfate, filtered and concentrated. The crude product was then purified by column chromatography (Hexanes: Ethyl Acetate 1:0 to 7:3) to give the pure product as an orange solid in 35% yield (330 mg). $^1H$ NMR (500 MHz, $CDCl_3$) δ 9.01 (s, 1H), 8.11 – 7.95 (m, 2H), 7.81 (d, *J* = 16.2 Hz, 1H), 7.74 – 7.70 (m, 1H), 7.66 – 7.63 (m, 1H), 7.56 (d, *J* = 8.8 Hz, 2H), 7.18 (d, *J* = 16.2 Hz, 1H), 6.74 (d, *J* = 8.7 Hz, 2H), 3.03 (s, 7H). $^{13}C$ NMR (125 MHz, $CDCl_3$) δ 151.69, 151.22, 144.68, 142.61, 141.22, 137.06, 130.25, 129.19, 129.12, 128.92, 128.64, 124.18, 120.59, 112.25, 40.41. HRMS (ESI+): *m/z*. 276.14938 [$C_{18}H_{18}N_3$ $(M+H)^+$, calcd 276.14952].

**FLA**

An aqueous solution of NaOH (48.0 g, 120 mmol) in 80 mL $H_2O$, was added slowly to a solution of 2-hydroxyacetophenone (4.80 g, 35.2 mmol) in a solution of MeOH (100 mL). Upon cooling to room temperature, 4-dimethylaminobenzaldehyde (5.97 g, 33.3 mmol) was added to the reaction mixture and stirred at room temperature overnight. Aqueous solution of NaOH (9.6 g, 240 mmol) in 48 mL $H_2O$ was then added, followed by dropwise addition of 30% $H_2O_2$ solution (40 mL) to the reaction mixture and stirred at room temperature overnight. Upon completion of the reaction, the precipitates were filtered, washed with water and dried to give the pure product as a yellow solid in 45% yield (4.20 g). $^1H$ NMR (400 MHz, $CDCl_3$) δ 8.23 (dd, J = 8.0, 1.5 Hz), 8.19 (d, J = 9.2 Hz), 7.65 (ddd, J = 8.6, 7.0, 1.7 Hz), 7.55 (d, J = 8.0 Hz), 7.38 (ddd, J = 8.1, 7.1, 1.1 Hz), 6.94 (s), 6.80 (d, J = 9.2 Hz), 3.07 (s). $^{13}C$ NMR (100

MHz, $CDCl_3$) δ 172.60, 155.18, 151.42, 146.71, 137.04, 132.91, 129.29, 125.32, 124.24, 120.96, 118.26, 118.11, 111.58, 40.17. HRMS (ESI+): *m*/*z*. 282.11255 [$C_{17}H_{16}NO_3$ $(M+H)^+$, calcd 282.11247].

**FLAM**

**FLA** (1.50 g, 5.34 mmol) and $K_2CO_3$ (1.48 g, 10.7 mmol) were added to acetone (30 mL). To this mixture, iodomethane (2.26 g, 16.0 mmol) was added and the resulting reaction mixture was stirred at room temperature overnight. Upon completion, the reaction mixture was extracted with DCM (3x) and washed with water (3x), dried over magnesium sulfate, filtered and concentrated. The crude product was then purified by column chromatography (Hexanes: DCM, 1:0 to 9:1) to give the pure product as a yellow solid in 74% yield (1.16 g). $^1H$ NMR (400 MHz, $CHCl_3$) δ 8.25 (dd, *J* = 8.0, 1.6 Hz), 8.16 – 8.04 (m), 7.63 (ddd, *J* = 8.6, 7.1, 1.6 Hz), 7.53 – 7.46 (m), 7.39 – 7.31 (m), 6.84 – 6.68 (m), 3.88 (s), 3.07 (s). $^{13}C$ NMR (100 MHz, $CHCl_3$) δ 174.77, 156.64, 155.19, 151.82, 140.26, 132.99, 130.02, 125.80, 124.40, 117.86, 111.45, 59.81, 40.17. HRMS (ESI+): *m*/*z*. 296.12818 [$C_{18}H_{18}NO_3$ $(M+H)^+$, calcd 296.12812].

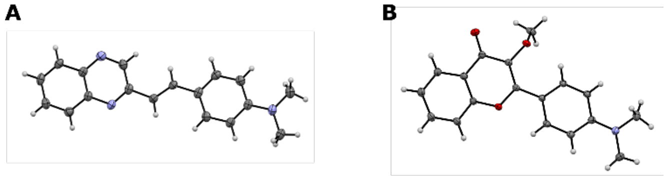


**QX-DMA** **FLAM**

**Figure S1**: Crystal structure of environment responsive probes. (A) Structure of **QX-DMA** with R-factor of 5.20%. (B) Structure of **FLAM** with R-factor of 6.48%.

**Table S1**: Crystal data and structure refinement for **QX-DMA**.

| | |
|---|---|
| Identification code | **QX-DMA** |
| Empirical formula | $C_{18}H_{17}N_3$ |
| Formula weight | 275.34 |
| Temperature/K | 100.00(10) |
| Crystal system | monoclinic |
| Space group | $P2_1/n$ |
| a/Å | 15.0623(7) |
| b/Å | 6.2873(2) |
| c/Å | 15.1919(6) |
| α/° | 90 |
| β/° | 99.821(4) |
| γ/° | 90 |
| Volume/$Å^3$ | 1417.61(10) |
| Z | 4 |
| $\rho_{calc}g/cm^3$ | 1.290 |
| $\mu/mm^{-1}$ | 0.606 |

| | |
|---|---|
| F(000) | 584.0 |
| Crystal size/mm$^3$ | 0.201 × 0.041 × 0.021 |
| Radiation | Cu Kα (λ = 1.54184) |
| 2Θ range for data collection/° | 7.64 to 155.844 |
| Index ranges | -19 ≤ h ≤ 18, -6 ≤ k ≤ 7, -19 ≤ l ≤ 18 |
| Reflections collected | 15559 |
| Independent reflections | 2967 [$R_{int}$ = 0.0529, $R_{sigma}$ = 0.0397] |
| Data/restraints/parameters | 2967/39/211 |
| Goodness-of-fit on F$^2$ | 1.079 |
| Final R indexes [I>=2σ (I)] | $R_1$ = 0.0520, $wR_2$ = 0.1330 |
| Final R indexes [all data] | $R_1$ = 0.0647, $wR_2$ = 0.1406 |
| Largest diff. peak/hole / e Å$^{-3}$ | 0.25/-0.25 |

**Table S2**: Crystal data and structure refinement for **FLAM**.

| | |
|---|---|
| Identification code | **FLAM** |
| Empirical formula | $C_{18}H_{17}NO_3$ |
| Formula weight | 295.32 |
| Temperature/K | 100.0(2) |
| Crystal system | triclinic |
| Space group | P-1 |
| a/Å | 9.7916(2) |
| b/Å | 12.1263(2) |
| c/Å | 12.9449(2) |
| α/° | 92.250(2) |
| β/° | 94.3000(10) |
| γ/° | 108.529(2) |
| Volume/Å$^3$ | 1450.01(5) |
| Z | 4 |
| $\rho_{calc}$g/cm$^3$ | 1.353 |
| μ/mm$^{-1}$ | 0.092 |
| F(000) | 624.0 |
| Crystal size/mm$^3$ | 0.201 × 0.041 × 0.021 |
| Radiation | Mo Kα (λ = 0.71073) |
| 2Θ range for data collection/° | 4.91 to 115.992 |
| Index ranges | -20 ≤ h ≤ 23, -28 ≤ k ≤ 27, -30 ≤ l ≤ 30 |
| Reflections collected | 103131 |
| Independent reflections | 39581 [$R_{int}$ = 0.0769, $R_{sigma}$ = 0.1074] |
| Data/restraints/parameters | 39581/0/403 |
| Goodness-of-fit on F$^2$ | 0.968 |
| Final R indexes [I>=2σ (I)] | $R_1$ = 0.0648, $wR_2$ = 0.1588 |
| Final R indexes [all data] | $R_1$ = 0.1649, $wR_2$ = 0.2065 |
| Largest diff. peak/hole / e Å$^{-3}$ | 0.62/-0.34 |

#### 3 Photophysical Studies

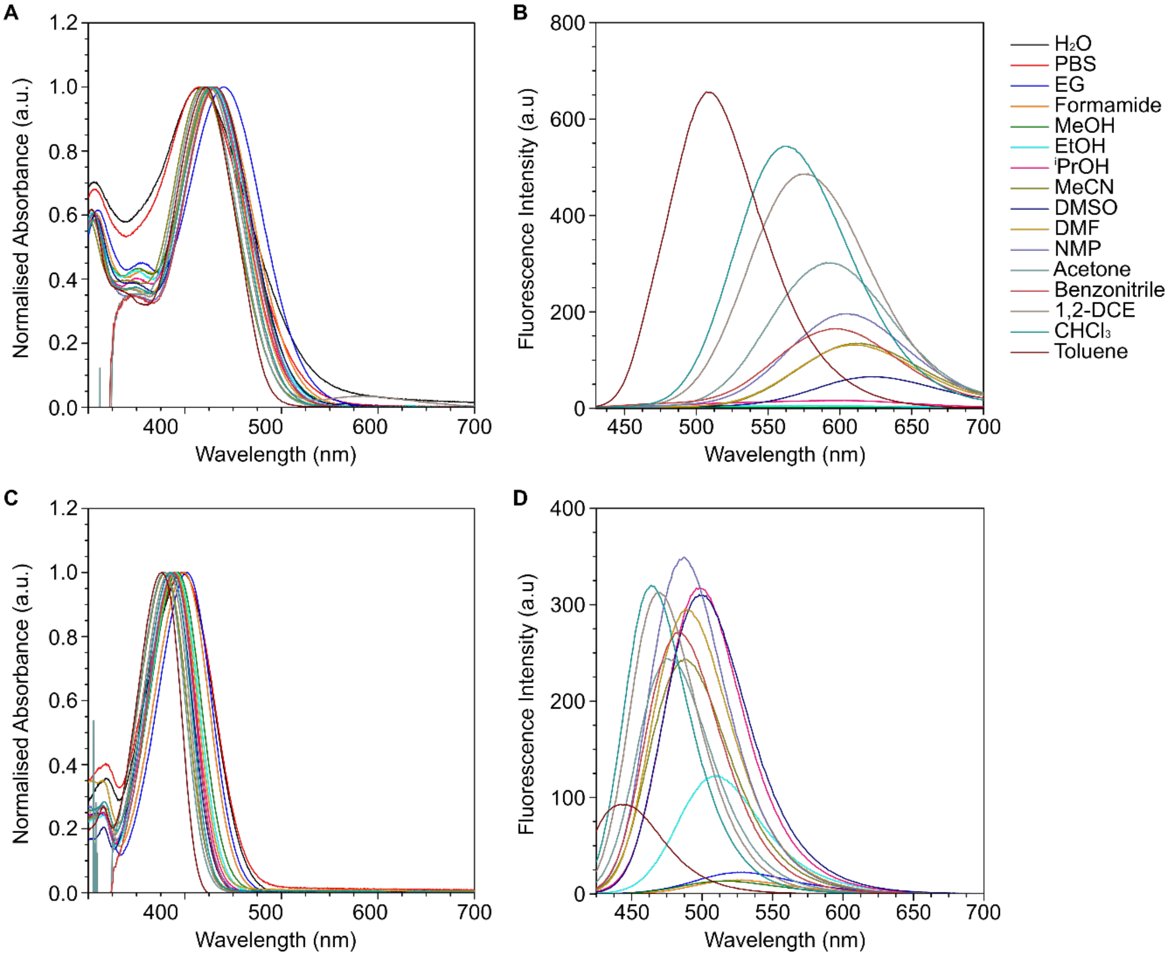


**Figure S2**: Comparison of photophysical characterisation of solvatochromic behaviour of dyes. (A) Normalised absorbance spectra of **QX-DMA**. (B) Fluorescence emission spectra of **QX-DMA**. (C) Normalised absorbance spectra of **FLAM**. (B) Fluorescence emission spectra of **FLAM**. 405 nm excitation wavelength was used for fluorescence emission spectra measurements. PBS was $Na_2HPO_4$ at 20 mM concentration. 10 µM dye concentration was used for all measurements. 1,2-DCE: 1,2-dichloroethan; iPrOH: isopropanol; EG: ethylene glycol.

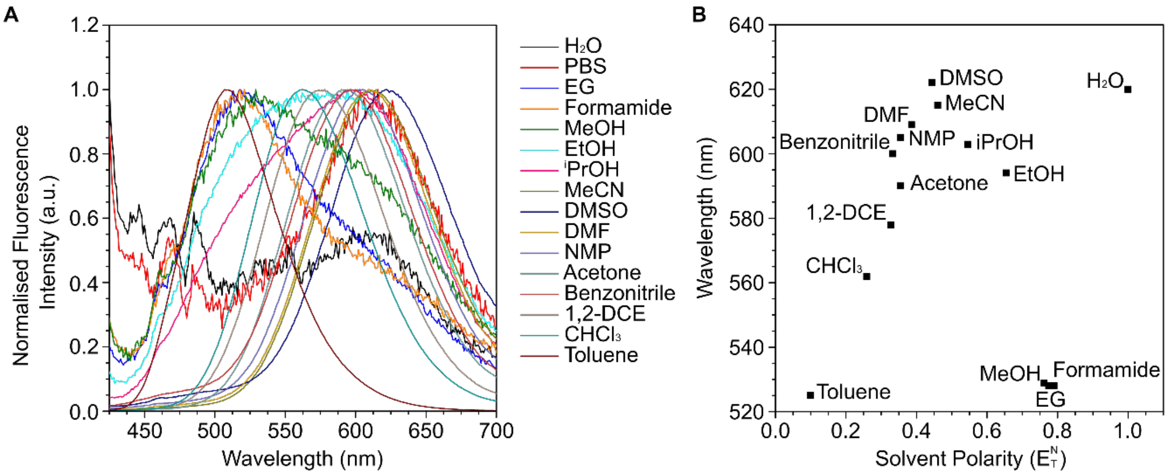


**Figure S3**: Normalised fluorescence emission spectra and relationship with solvent polarity ($E_T^N$) for **QX-DMA** and **FLAM**. (A, C) Normalised fluorescence emission spectra for **QX-DMA** and **FLAM** respectively, measured in a range of polarities. (B, D) Relationship of measured fluorescence emission maxima wavelength and solvent polarity for **QX-DMA** and **FLAM** respectively. Solvent polarity was based on reported values, with PBS excluded from linear plot (see B, D).

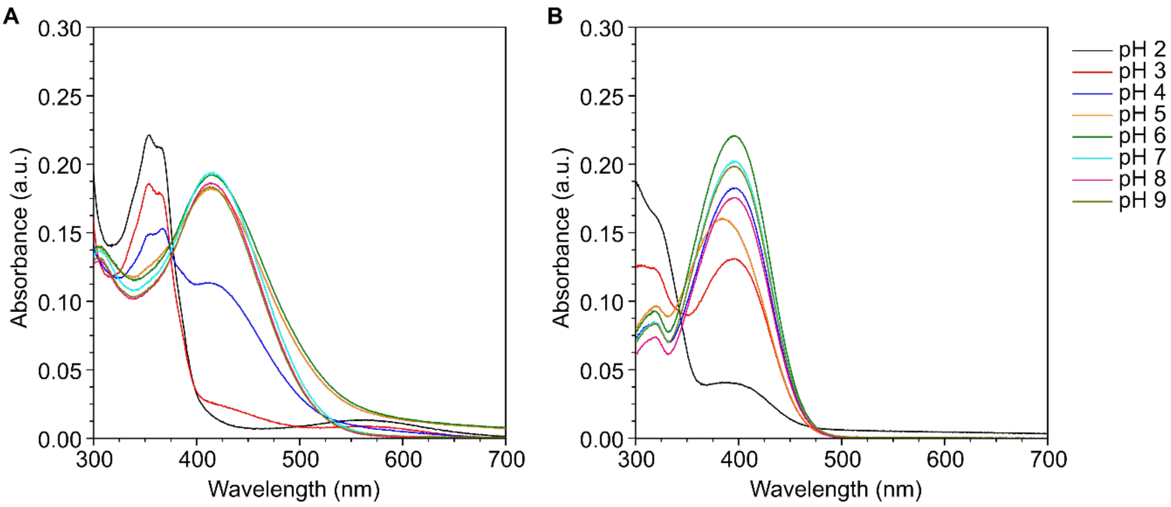


**Figure S4**: Measurement of dyes in physiologically relevant pH. (A) Absorbance of **QX-DMA**. (B) Absorbance of **FLAM**. 10 µM dye concentration was used for all measurements.

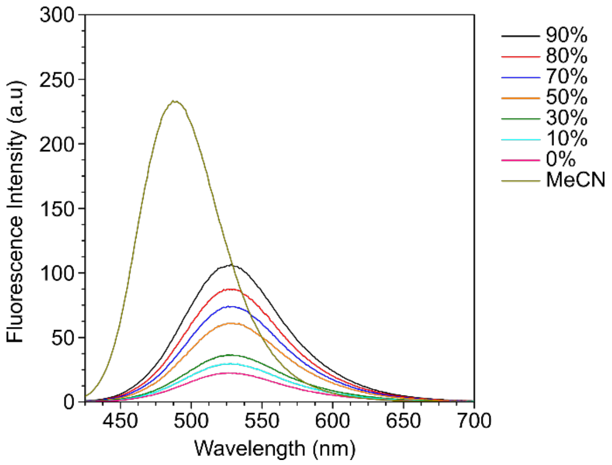


**Figure S3**: Viscosity experiment with varying ratios of ethylene glycol and glycerol for **FLAM**. 405 nm excitation wavelength was used. 10 μM dye concentration was used for all measurements.

#### 4 Biological Studies

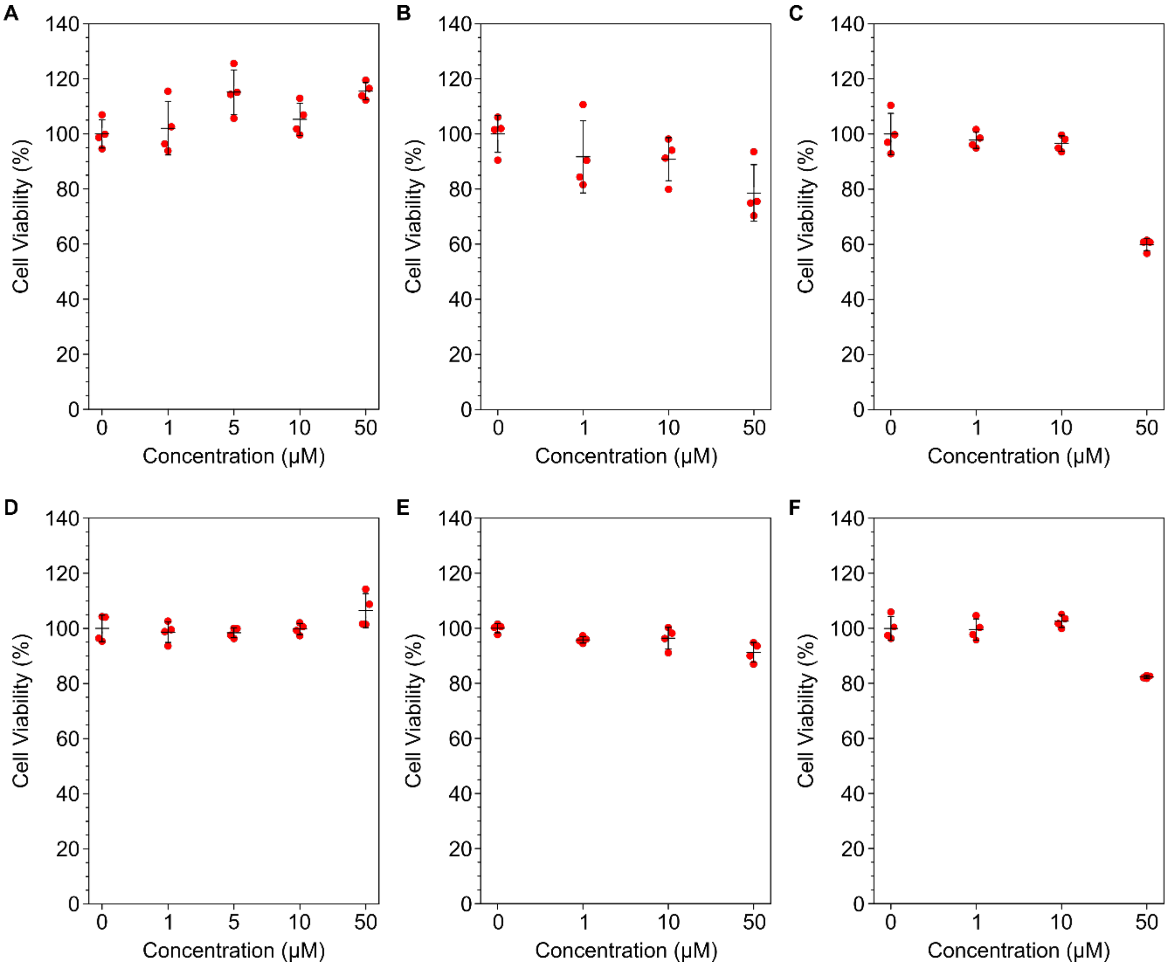


**Figure S4**: Cytotoxicity assay for **FLAM** in HeLa and A549 cells. (A, B, C) HeLa cell viability after 0.5, 24, 48 h dye incubation respectively. (D, E, F) A549 cell viability after 0.5, 24, 48 h dye incubation respectively. Resazurin (alamarBlue™) cell viability assay was recorded by a plate reader. n = 4 biological replicates; mean ± s.d.

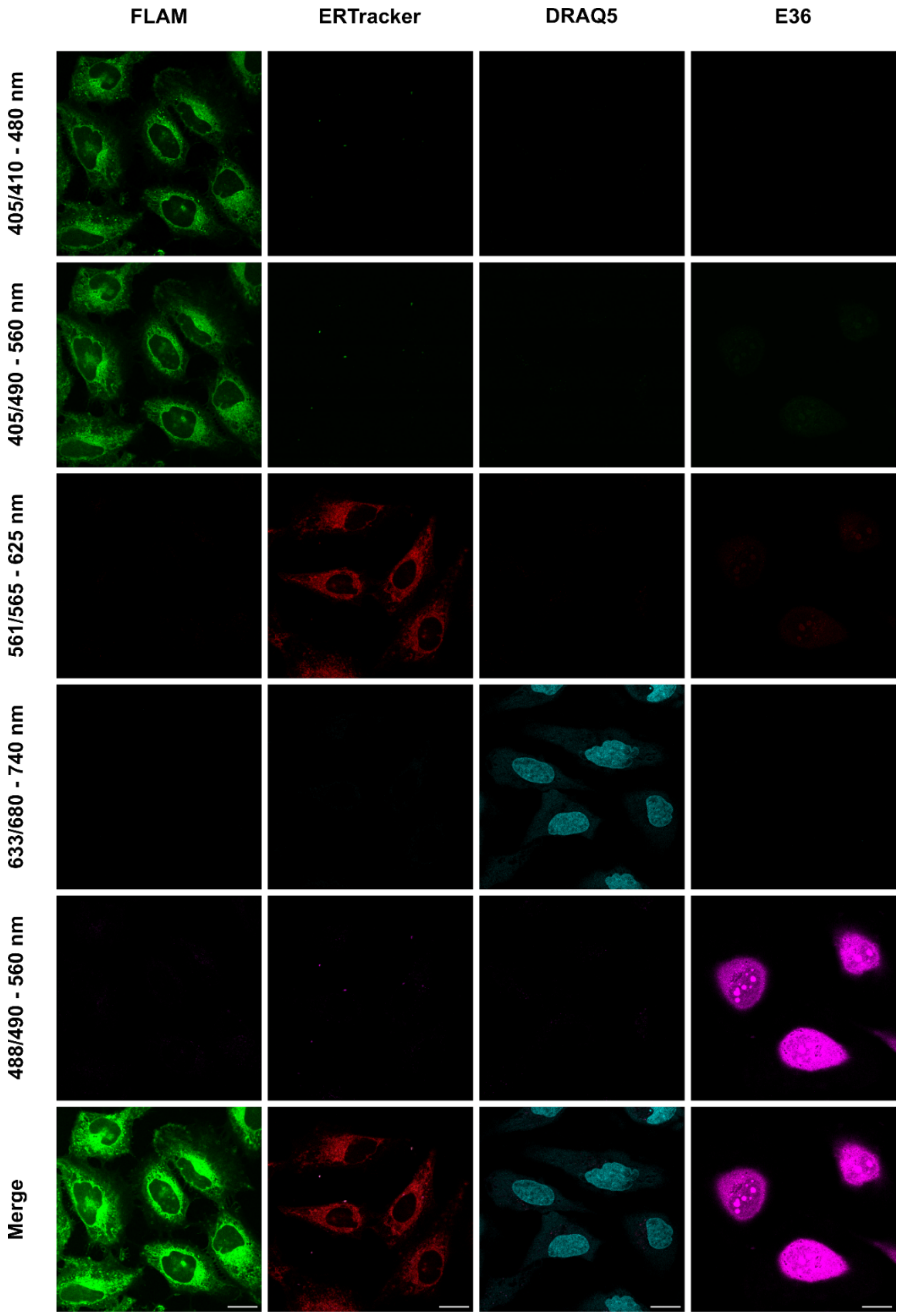


**Figure S5**: Control microscopy experiments for dye crosstalk in HeLa cells. ERTracker™ Red, DRAQ5™ and E36 stains were used to visualise endoplasmic reticulum, nucleus (DNA) and RNA respectively. Cells were stained with 5 μM **FLAM**, 1 μM DRAQ5™ and E36, 500 nM ERTracker™ Red. Scale bar, 20 μM.

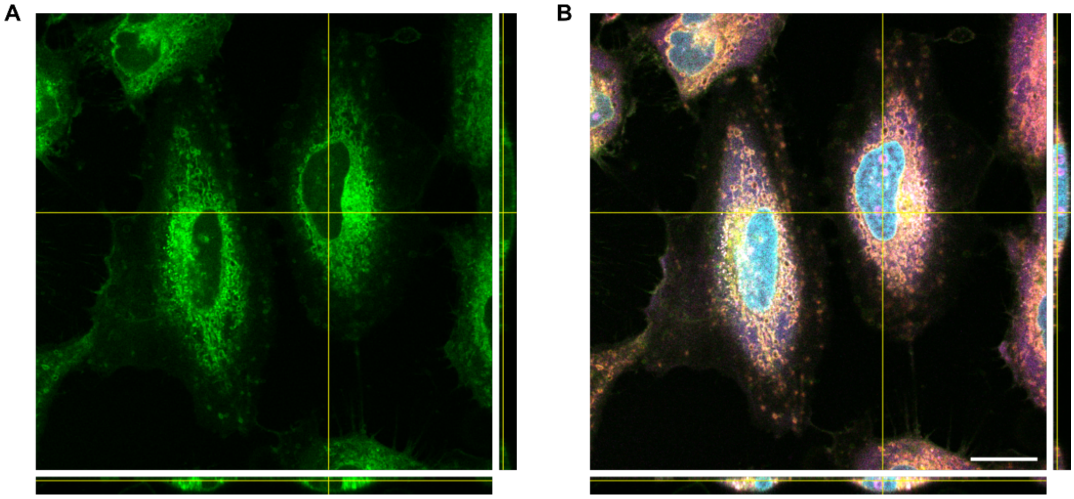


**Figure S6**: Z-stack cell localisation microscopy experiments for stained HeLa cells. (A) Orthoslice visualisation of **FLAM**. (B) Merged orthoslice visualisation of **FLAM** and co-staining dyes. ERTracker™ Red. DRAQ5™ and E36 stains were used to visualise endoplasmic reticulum, nucleus (DNA) and RNA respectively. Cells were stained with 5 μM **FLAM** (green), 1 μM DRAQ5™ (cyan) and E36 (magenta), and 500 nM ERTracker™ Red (red). 405/490 – 560 nm channel of **FLAM** shown. Scale bar, 20 μm.

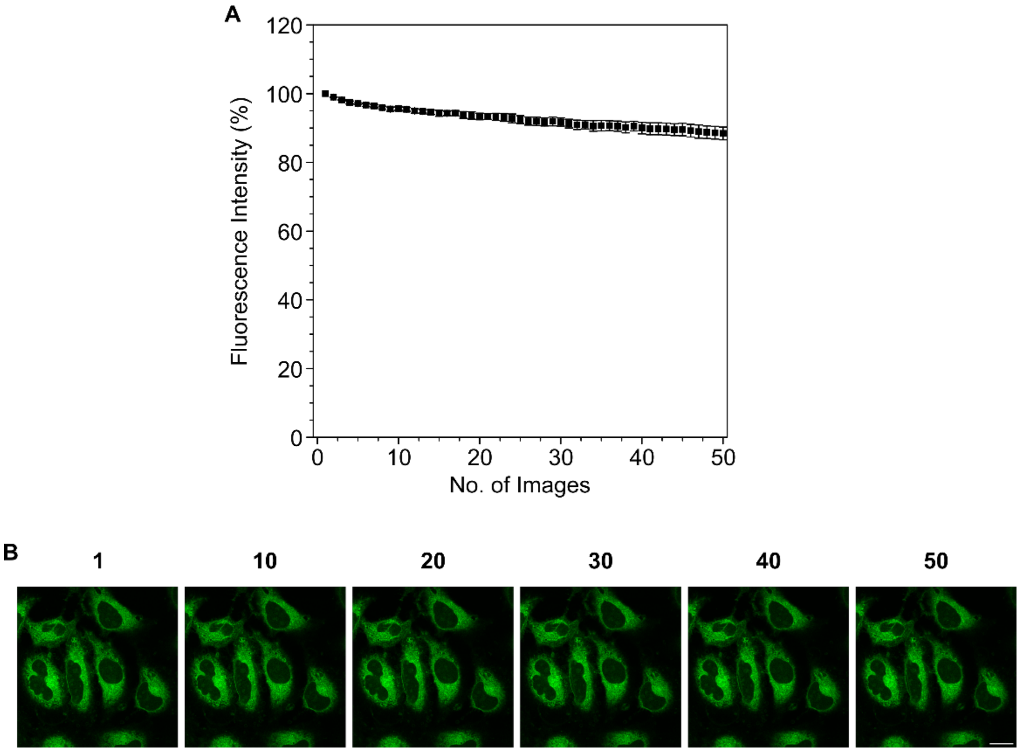


**Figure S7**: Photostability microscopy experiments for **FLAM** stained HeLa cells. (A) Fluorescence intensity of **FLAM**. (B) Images of **FLAM** stained cells at selected points. Cells were stained with 5 μM **FLAM** (green). 405/ 490 – 560 nm channel of **FLAM** shown. Scale bar, 20 μm. n = 5 cells analysed; mean ± s.d.

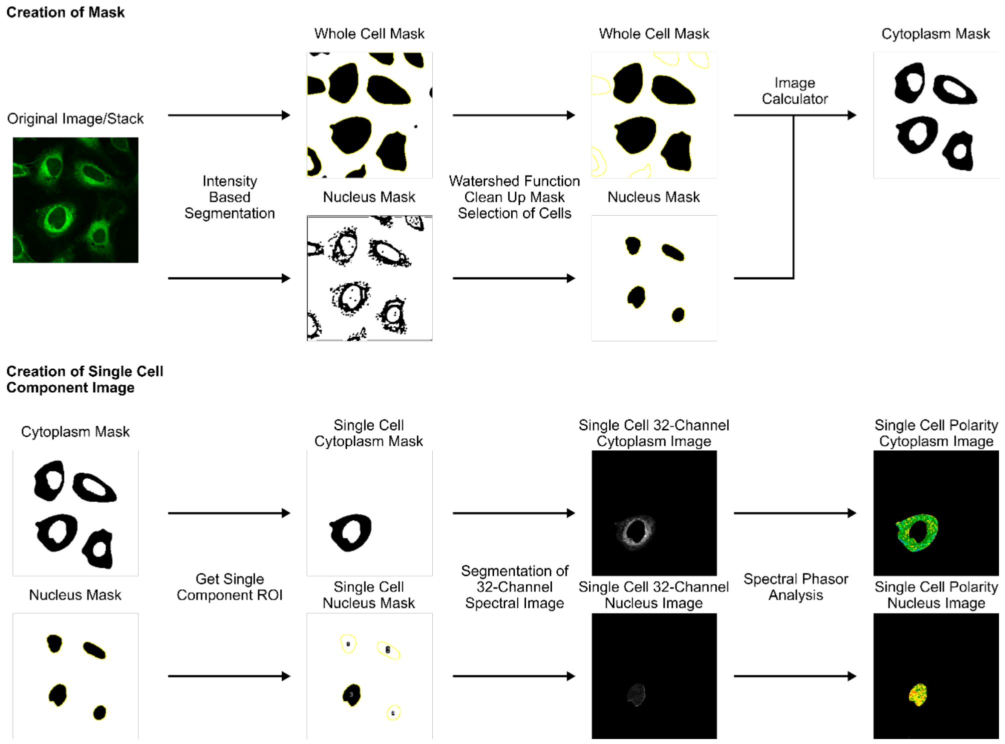


**Figure S8**: Flow chart for segmentation of cell components and 32-channel spectral images by image processing scripts.

### 5 $^{1}$H and $^{13}$C NMR Spectra for compounds

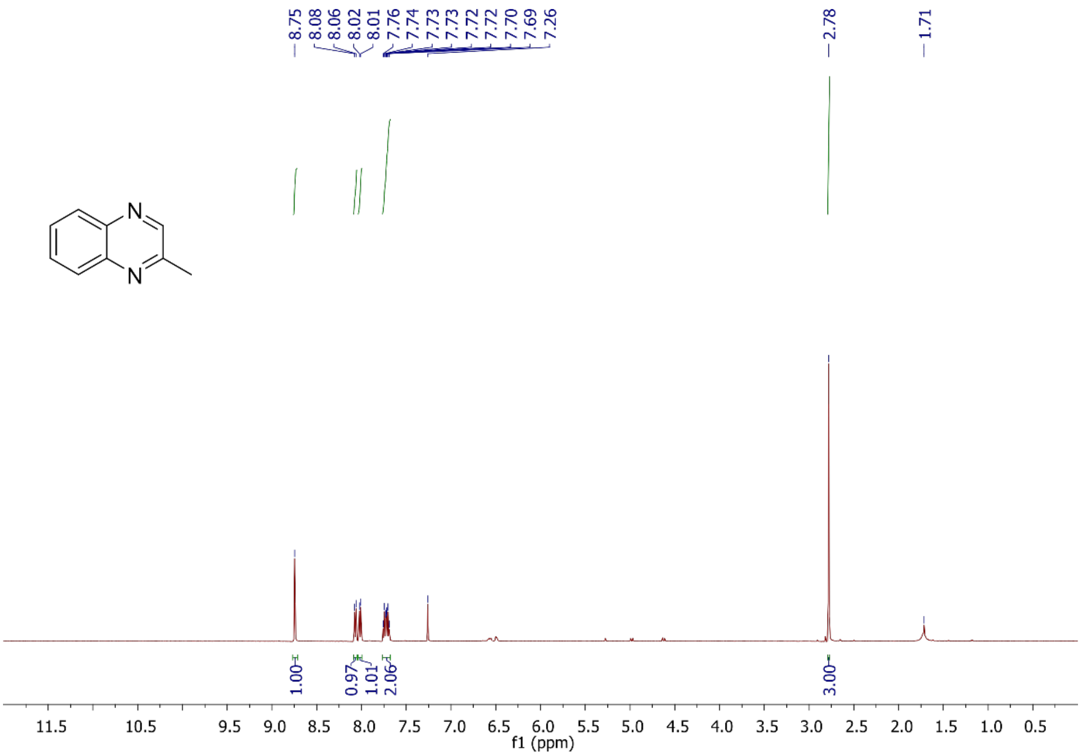


Figure S9: $^{1}$H NMR of **QX** in $CDCl_3$.

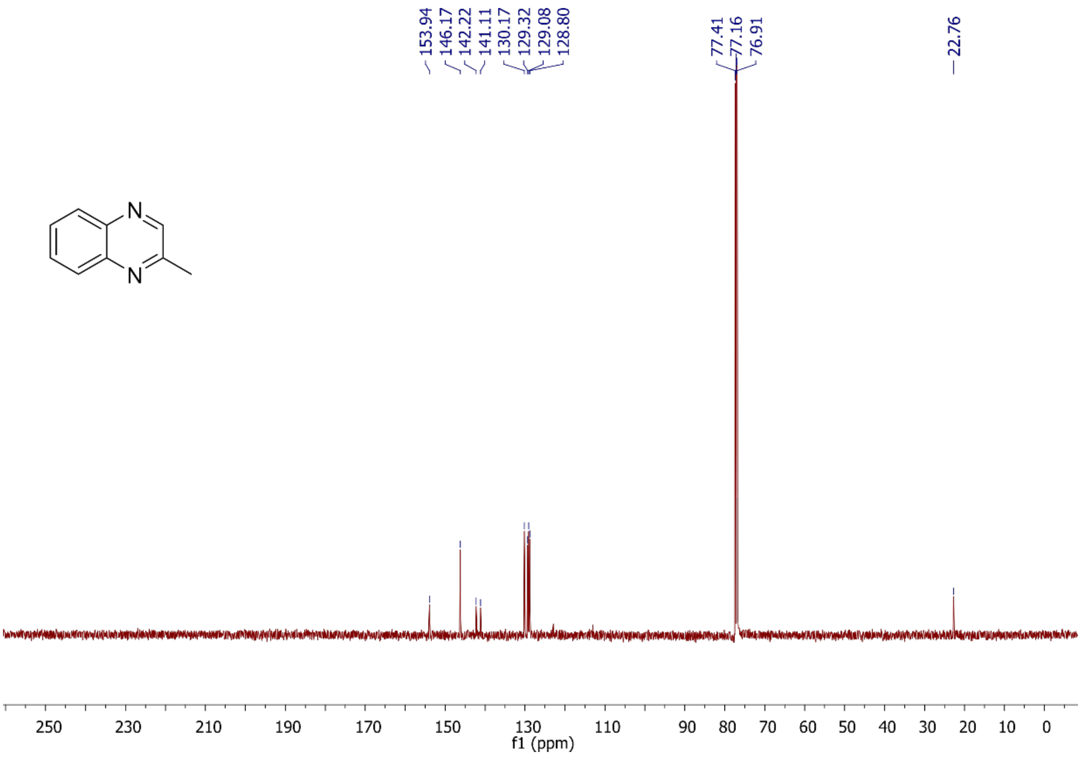


Figure S10: $^{13}$C NMR of **QX** in $CDCl_3$.

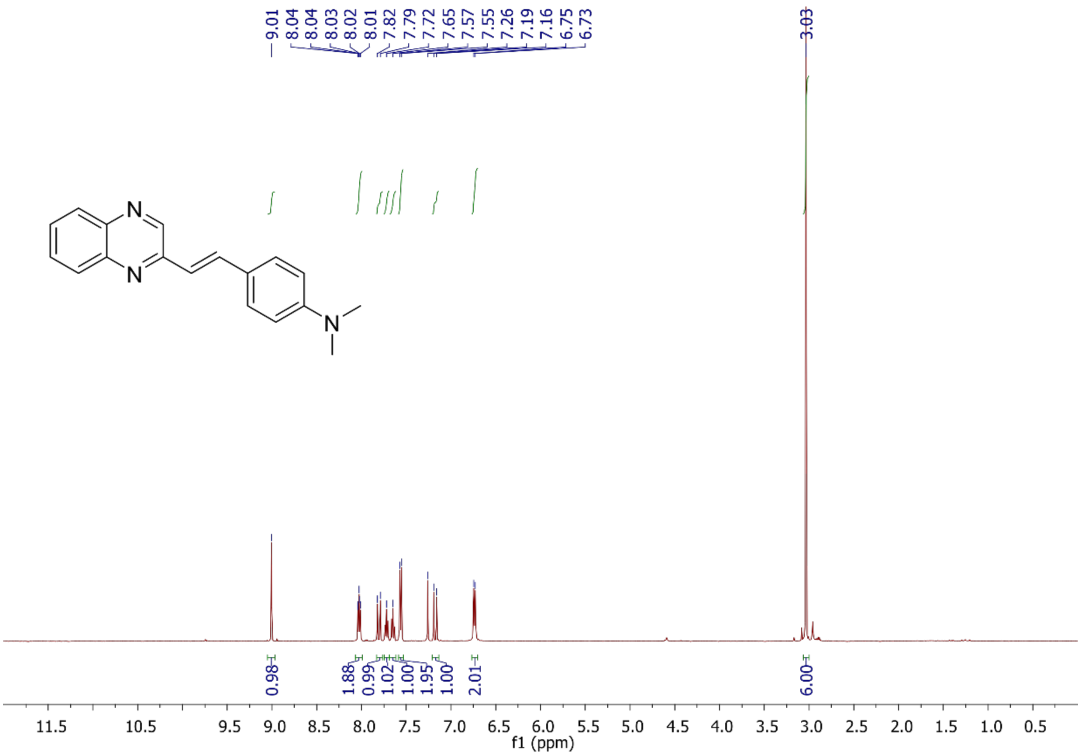


Figure S11: $^{1}$H NMR of **QX-DMA** in $CDCl_3$.

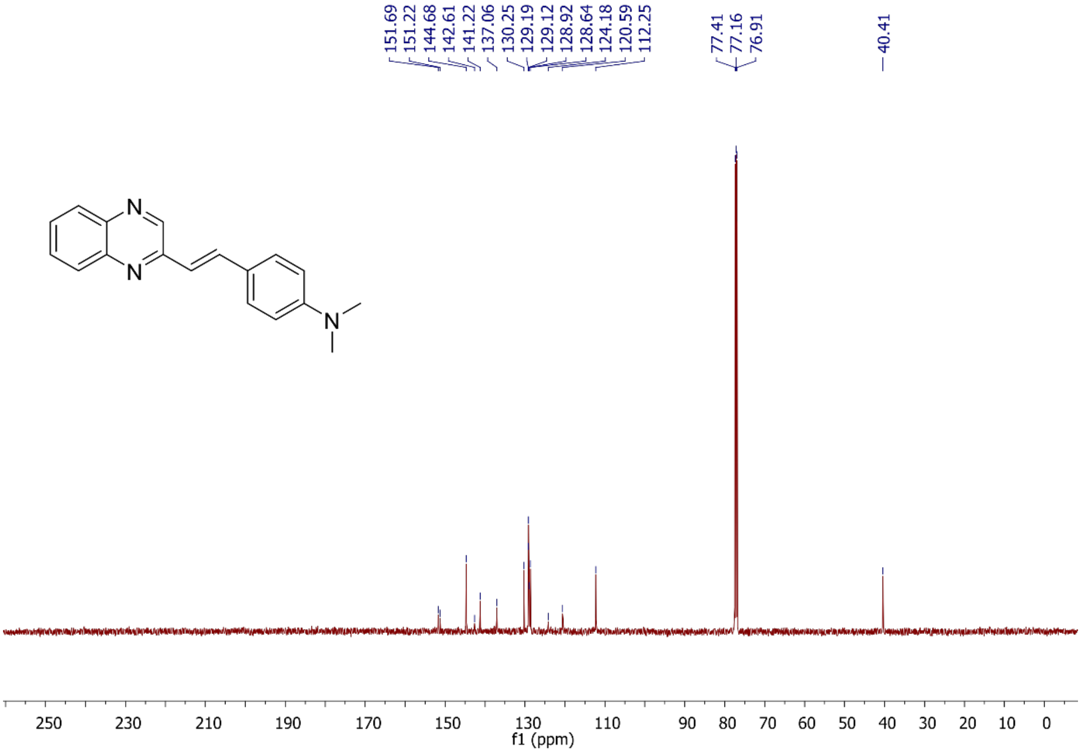


Figure S12: $^{13}$C NMR of **QX-DMA** in $CDCl_3$.

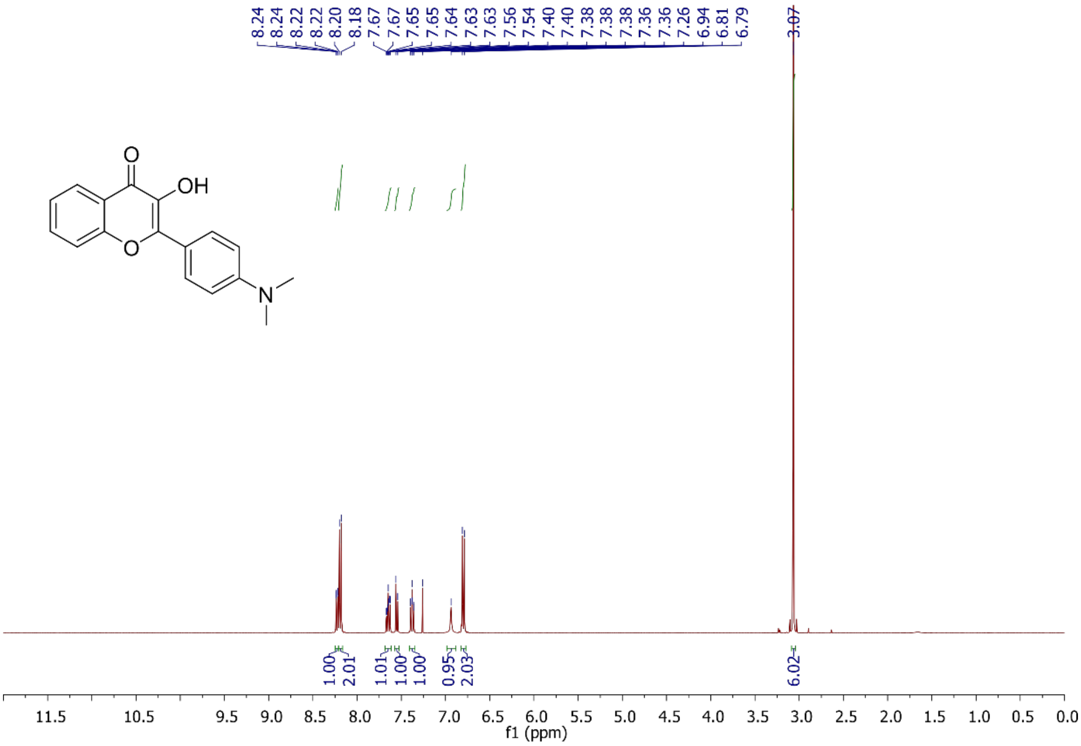


Figure S13: $^{1}H$ NMR of **FLA** in $CDCl_3$.

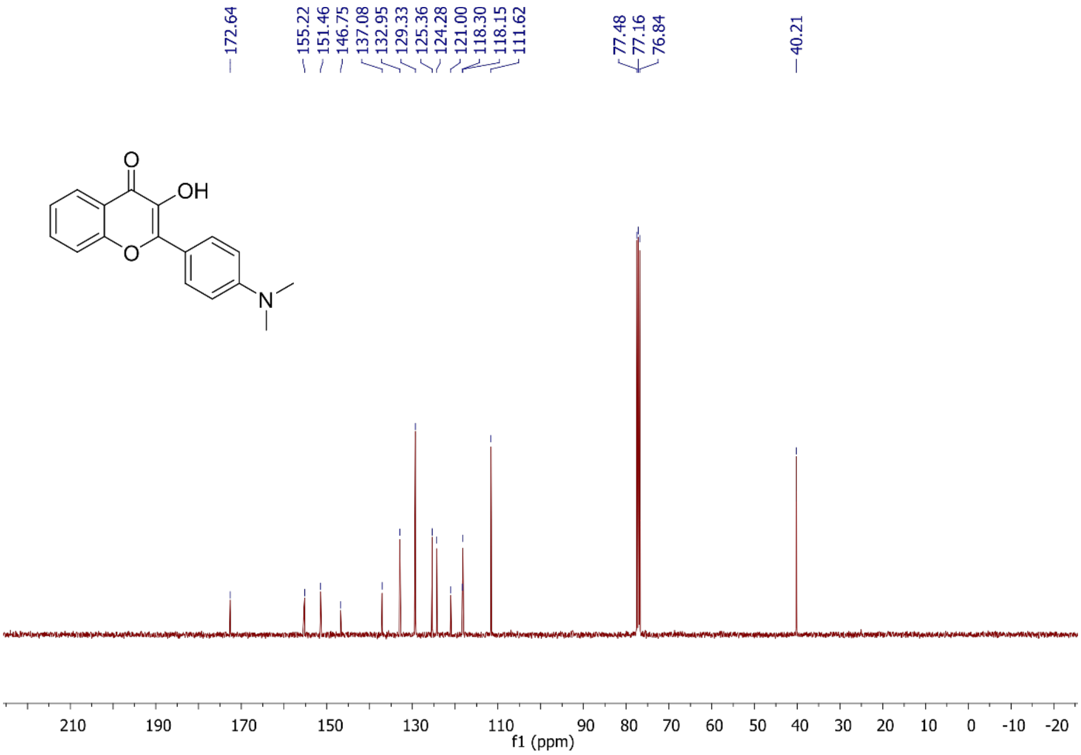


Figure S14: $^{13}C$ NMR of **FLA** in $CDCl_3$.

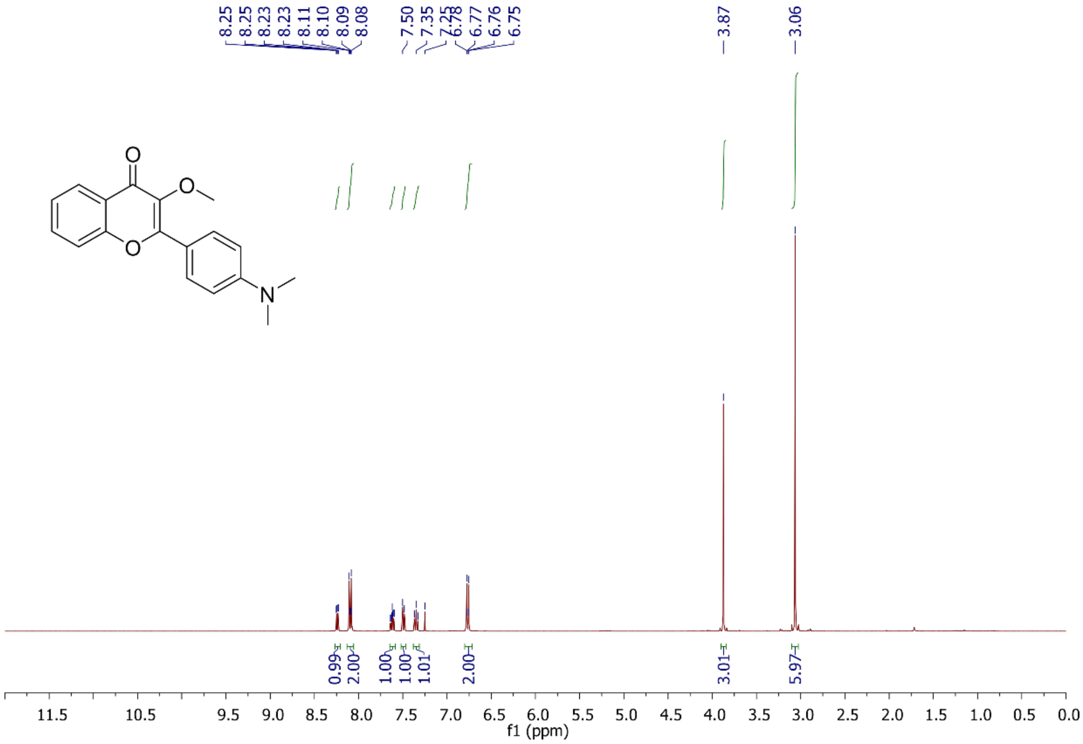


Figure S15: [1]H NMR of **FLAM** in $CDCl_3$.

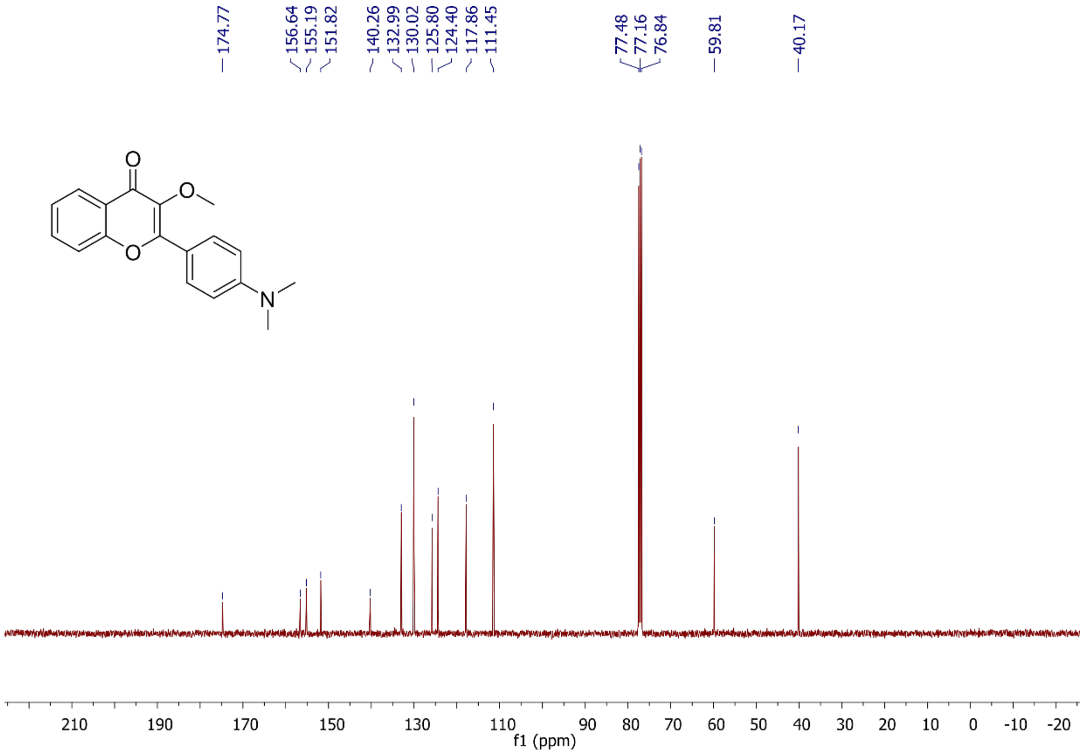


Figure S16: [13]C NMR of **FLAM** in $CDCl_3$.
